# Naturalistic language comprehension is supported by alpha and beta oscillations linked to domain-general inhibition and reactivation

**DOI:** 10.1101/2022.08.05.502909

**Authors:** Ioanna Zioga, Hugo Weissbart, Ashley G. Lewis, Saskia Haegens, Andrea E. Martin

**Author notes:** **Corresponding author:** Ioanna Zioga.

## Abstract

Brain oscillations are prevalent in all species and are involved in numerous perceptual operations. Alpha oscillations are thought to facilitate processing through the inhibition of task-irrelevant networks, while beta oscillations are linked to the reactivation of content representations. Can the proposed functional role of alpha and beta oscillations be generalized from low-level operations to higher-level cognitive processes? Here we address this question focusing on naturalistic spoken language processing. Twenty-two (18 female) Dutch native speakers listened to stories in spoken Dutch and French while magnetoencephalography (MEG) was recorded. We used dependency parsing to identify three dependency states at each word, as the number of (1) newly opened dependencies, (2) dependencies that remained open, and (3) resolved dependencies. We then constructed linear forward models to predict alpha and beta power from the dependency features. Results showed that dependency features predict alpha and beta power in language-related regions beyond low-level linguistic features. Left temporal, fundamental language regions are involved in language comprehension in the alpha band, while frontal and parietal, higher-order language regions, and motor regions are mostly involved in the beta band. Critically, alpha and beta band dynamics seem to subserve language comprehension tapping into syntactic structure building and semantic composition by providing low-level mechanistic operations for inhibition and reactivation processes. Overall, this study sheds light on the role of alpha and beta oscillations during naturalistic language processing, providing evidence for the generalizability of these dynamics from perceptual to complex linguistic processes.

**Significance Statement:** Prior research identified the functional role of alpha and beta oscillations in basic perceptual and motor functions. However, it remains unclear whether their proposed role can be generalized to higher-level processes during language comprehension. Here, we found that high-level syntactic features predict alpha and beta power in language-related regions beyond low-level linguistic features when listening to comprehensible naturalistic speech. Our work contributes to the debate about whether the functional roles of brain oscillations are domain-general or depend on the task at hand. We offer experimental findings that integrate a neuroscientific framework on the role of brain oscillations as “building blocks” with language comprehension as a compositional process, and novel evidence regarding the encoding of higher-level syntactic operations in the brain. This supports the view of a domain-general role of cortical oscillations across the hierarchy of cognitive functions, from low-level sensory operations to complex linguistic processes.

## 1. Introduction

Out of the many neural phenomena that exist, brain oscillations are prevalent in all species and are involved in numerous perceptual operations. Are brain oscillations the “building blocks” of cognitive function, from low-level sensory to higher-level processes? Extensive prior research focused on the role of alpha and beta oscillations in basic perceptual and motor functions. Here, we asked whether their proposed role in low-level operations generalizes to higher-level cognitive functions, in particular language comprehension. We adopted a forward-modelling approach predicting brain responses from high-level, syntactic features to test the role of alpha and beta oscillations during naturalistic language processing.

Alpha oscillations (8-12 Hz) are thought to reflect a mechanism of active inhibition, which fine-tunes sensory processing by guiding attention and suppressing distracting input (Jensen & Mazaheri, 2010; Klimesch et al., 2007). Alpha power increases with working memory load during retention, reflecting inhibition of task-irrelevant regions (Jensen et al., 2002; Pan et al., 2018; Scheeringa et al., 2009; Tuladhar et al., 2007), and is associated with behavioural performance (Haegens et al., 2010, 2011, 2012). Traditionally, beta oscillations (15-30 Hz) are considered a motor rhythm (Kilavik et al., 2013), associated with top-down processing and long-distance network communication (Varela et al., 2001). Beta power increases during information retention, attributed to active maintenance of the current cognitive set (Engel & Fries, 2010). Recently, Spitzer and Haegens (2017) proposed that beta oscillations support the reactivation of content representations, via the transitioning of a latent item into active working memory (Rose et al., 2016). To date, there is extensive research investigating the oscillatory correlates of language processing (Bastiaansen et al., 2010; Brennan & Martin, 2020; Coopmans & Cohn, 2022; Hauswald et al., 2020; Kaufeld et al., 2020; Kösem et al., 2016; Lewis et al., 2015; Meyer et al., 2020; Obleser & Weisz, 2012; van Bree et al., 2021; Weissbart et al., 2020; Zoefel & VanRullen, 2015), altogether demonstrating that a handful of basic oscillatory mechanisms are responsible for a plethora of linguistic operations (Prystauka & Lewis 2019). However, little is known yet as to whether alpha and beta oscillations fulfill the same function in language comprehension as in low-level sensory tasks.

Here, we operationalized high-level linguistic processing using attributes from dependency parsing, which describes syntactic sentence structure as relations between pairs of words (Mel’cuk, 1988; Tesnière, 2015). Dependencies are created when a non-unified word (“dependent constituent”) is encountered. Processing load increases with the number of dependencies being processed (Demberg & Keller, 2008; Vos et al., 2001). Dependencies are resolved once the linking word (“dependent”) is encountered, recruiting unification or integration processes (Hagoort, 2005; Kapteijns & Hintz, 2021; perceptual cue integration: Martin, 2016, 2020). Integration is thought to require reactivation of any dependent constituent (Foraker & McElree, 2011; Martin & McElree, 2008; McElree et al., 2003). As resolving linguistic dependencies is crucial for language comprehension, it can be argued that dependency resolution exemplifies higher-level cognitive operations. We therefore hypothesized that alpha and beta power would be modulated by dependencies, and used dependency parsing as a proxy for this processing.

Specifically, we compared MEG responses while participants listened to intelligible (Dutch) vs. unintelligible (French) spoken stories. We identified three states at each word: number of opened/remained open/resolved dependencies. We constructed forward models to predict alpha and beta power from these dependency features, controlling for low-level linguistic features. We predicted that Dutch compared to French stories would elicit stronger modulations of alpha and beta power in typical language processing brain regions. Additionally, we hypothesized that our dependency features would predict alpha and beta power in language processing-related regions beyond low-level linguistic features. We further hypothesized that the opening of dependencies would be associated with alpha in/decreases at task-ir/relevant areas linked to inhibition processes, while “maintenance” of dependencies would be associated with beta power increases, attributed to anticipatory and active ongoing processes. Finally, we predicted that beta power would increase during dependency resolution, related to content reactivation.

## 2. Materials and Methods

### 2.1. Participants

Twenty-two adults (18 female) aged between 18 and 63 years old (*M* ± *SD* age of 34 ± 15 years) took part in the experiment. Pre-screening required that participants were monolingual Dutch native speakers, right-handed, without hearing problems, reading problems or epilepsy. All participants self-reported zero use and minimal understanding of French at the level of isolated words but not whole sentences before taking part in the experiment. Prior to the experiment, participants were provided with written and verbal information about the MEG system and safety regulations and gave written informed consent. They received monetary reimbursement after participation. The study falls under the general ethics approval (CMO 2014/288 “Imaging Human Cognition”) in accordance with the Declaration of Helsinki.

### 2.2. Stories

In order to tap into language comprehension, we compared brain responses recorded with MEG while participants listened to intelligible (Dutch, mother tongue) vs. unintelligible (French, unfamiliar language) spoken stories. Stories in French were selected as a control to confirm that our effects are due to comprehension and not acoustic properties of speech, with which participants would be familiar. Dutch native speakers were not able to understand the French narratives, despite their familiarity with the acoustic properties and some common words in French. French thus constituted a stronger control than a language with which participants would be completely unfamiliar. Critically, compared to traditional studies using artificial word or sentence stimuli, the use of natural speech in pre-recorded stories allowed for a more ecologically-valid approach as (1) the natural prosody of the voice recording guides comprehension via auditory cues, (2) processing requires constant effortful attention throughout, and (3) it lacks the brain responses induced by certain properties of artificial stimuli, such as abrupt voice modulations or unnatural syllable timing.

The following three Dutch (NL) stories were used: *Het Lelijke Jonge Eendje* by H.C. Andersen, *De Ransel, het Hoedje en het Hoorntje* and *De Gouden Vogel* by the Grimm brothers. All NL stories were spoken by female voices. The following three French (FR) stories were used: *L’eau de la vie* by the Grimm brothers (male voice), *L’ange* by H.C. Andersen (female voice), and an excerpt from *Le Canard Ballon* by E.A. Poe (female voice). The NL stories and the last FR story were retrieved from www.librivox.org, and the rest from www.litteratureaudio.com. In order to reduce fatigue, stories were split into parts of short duration (NL: 9 story parts, *M* ± *SD* 5.5 ± .6 min; FR: 4 story parts, duration 5.3 ± .7 min). Stories that were already less than 6 min were not split further. All audio files were normalized to an equal perceived loudness.

Five MCQs with four choices each were included after each story part (65 in total) to (1) ensure that participants paid attention to the spoken stories, and (2) confirm the lack of understanding of the French stories. A Dutch and a French native speaker composed the questions for the Dutch and French stories, respectively. All were content questions, e.g., *Who lives in the old house? A. An old man B. An old lady C. Nobody D. A family; What did the traveler take from the table? A. The tablecloth B. The bread C. The potatoes D. The wine*.

### 2.3. Procedure

Participants were seated in the MEG system in a dimly lit room. They were informed that they would listen to stories in Dutch and French during MEG recording. Further, they were instructed to pay attention to the stories as they would be prompted to answer MCQs after each story part. Responding to the MCQs was done by pressing four keys of a response box in a self-paced manner. Resting state MEG was recorded for 10 s before the onset of each story part but was not included in the analysis. The presentation order of the story parts was pseudorandomized across participants: NL and FR story parts were interleaved but care was taken so that their order remained intact (e.g., the second part of a story could be presented only if the first part of that same story was previously heard). The overall procedure lasted for approximately 1.5 hr. The experiment was programmed with custom MATLAB (MathWorks) scripts using Psychtoolbox (Brainard & Vision, 1997).

### 2.4. Data acquisition and MEG preprocessing

MEG data were recorded at a sampling rate of 1,200 Hz using a 275-channel axial gradiometer system (CTF MEG systems, VSM MedTech Ltd.) located in a magnetically shielded room. Eight sensors were excluded due to permanent malfunction, leaving a total of 267 usable sensors. Three fiducial localization coils were placed at the participant’s nasion and left and right ear canals to (1) allow for real-time monitoring of the participant’s head position and adjustment in between story parts if necessary, and (2) provide anatomical landmarks for offline coregistration of the MEG data with T1-weighted MRI images for source reconstruction. After completion of the task, the xyz coordinates of the three fiducial points as well as the participant’s head shape were digitized using a Polhemus 3D tracking device (Polhemus, Colchester, Vermont, United States). Furthermore, individual structural MRI scans were acquired in a 3T Siemens Magnetom Skyra MR scanner (Erlangen, Germany) using earplugs with a drop of vitamin E at the subject’s ear canals to facilitate subsequent alignment with the MEG data.

Continuous MEG data were downsampled to 100 Hz and epoched from the onset until the offset of each story part. Data from sensors with consistently poor signal quality, as observed by visual inspection, were removed and interpolated based on neighbouring sensors. Finally, independent component analysis was performed to correct for eye-blinks and heart-beat artifacts. Custom-written scripts in MATLAB and the FieldTrip toolbox (Oostenveld et al., 2011) were used for analysis of the MEG data.

### 2.5. Data analysis

#### 2.5.1. Behavioural data

To assess participants’ understanding of the stories, we calculated the percentage of correct responses in the MCQs separately for NL and FR stories. A paired *t*-test was used to compare the two conditions and a one-sample *t*-test to compare performance accuracy to chance level at 25%.

#### 2.5.2. Source reconstruction of MEG data

##### MRI preprocessing

First, coregistration of the MRI with the CTF and Polhemus fiducials was performed. Individual MRIs were normalized in MNI space and segmented. Realistic volume conduction models were created for each participant based on the single-shell model of their MRIs (Nolte, 2003). For each participant, 5798 dipole positions were defined with an 8-mm resolution.

##### Spatial filters

A spatial filter for the source reconstruction analysis was calculated for each participant. Covariance matrices were computed over single trials (13 Dutch and French story parts) and then averaged. Leadfields for all grid points, combined with the covariance matrices, were used to compute a spatial filter with the Linearly Constrained Minimum Variance (LCMV; Veen et al., 1997) method. The source orientation was fixed to the dipole orientation with the highest strength.

#### 2.5.3. Forward Models predicting alpha and beta power from linguistic features

We attempted to quantify higher-level operations during spoken language comprehension in response to the processing of dependencies that opened/ remained open/ were resolved at each word. We then constructed forward models predicting alpha and beta power from the dependency features controlling for low-level linguistic features (acoustic edges, word onset, and word frequency).

To investigate the relationship between linguistic features and alpha and beta power, we constructed a time series for each feature. Each word feature was time-aligned with the auditory stimulus using the forced-alignment function of the web-service MAUS (Kisler et al., 2017). In order to align the linguistic features with the auditory stimuli, a single impulse-like value representing the magnitude of the feature was assigned at the onset of each word (except for acoustic edges where the impulses could be at different time points, see below).

##### Dependency features

###### Dependency parsing

We operationalized high-level linguistic processing during spoken language comprehension using attributes from dependency parses; this is mainly motivated by the tradeoff between coverage of features and accessibility in parsing models, it is not a strong theoretical commitment to one parsing framework over another. Dependency grammars describe the syntactic structure of a sentence as a set of relations between two words (Mel’cuk, 1988; Tesnière, 2015). The links begin from the head and end on the dependent word and are assigned a label representing the type of dependency (e.g., subject, object, determinant, etc.). Each sentence has a root, usually the verb, which is the head of the entire structure (see the graph in Fig. 1A, top, for an example of dependency parsing). Dependency grammars often reveal non-adjacent, complex dependencies. Previous work has employed dependency structures as a measure of or proxy for syntactic complexity, as words that form dependencies often appear in non-adjacent positions (e.g., Wilson et al., 2020).

**Fig. 1.**
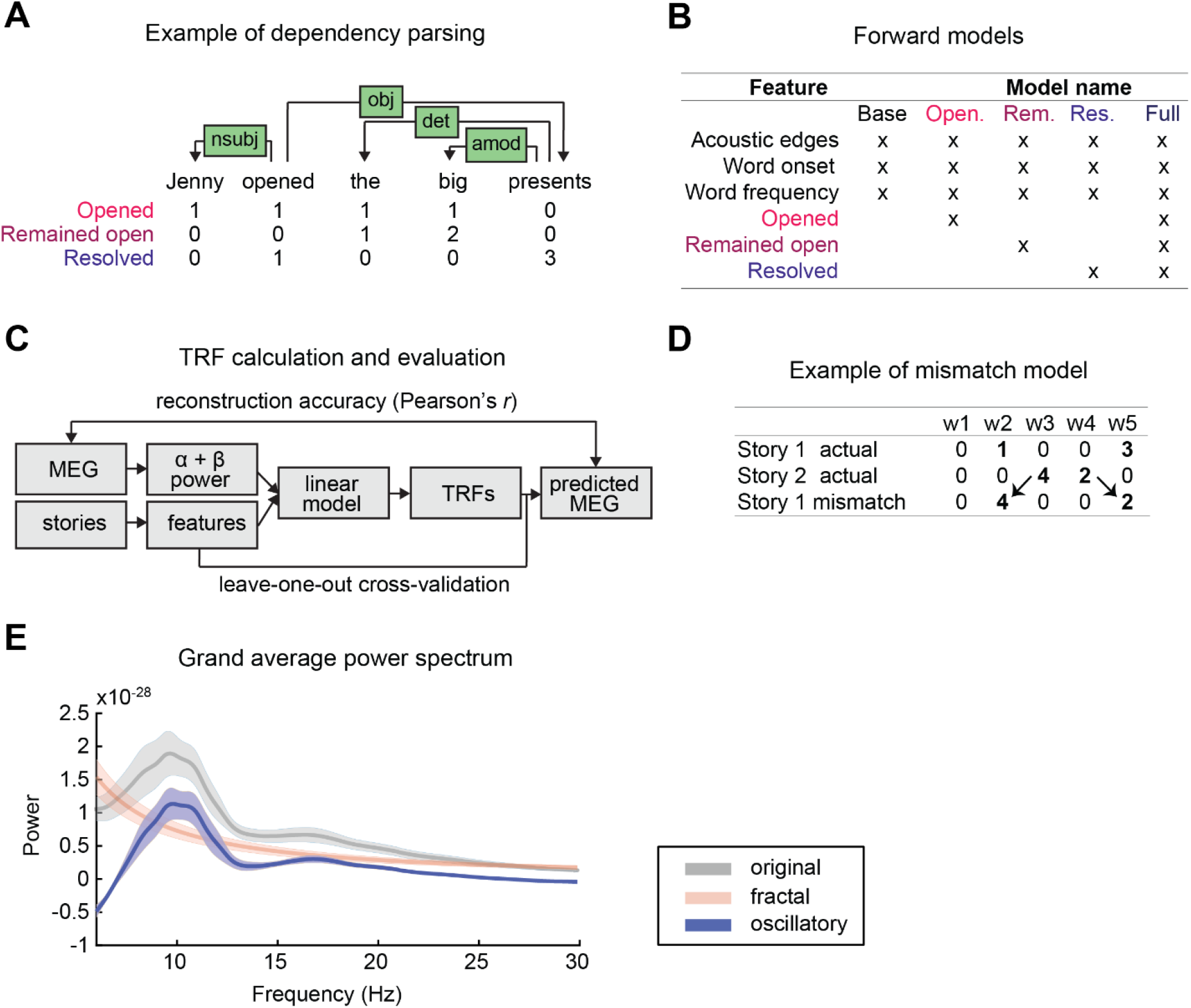
Methodological aspects of the Temporal Response Function (TRF) analysis using linguistic features as predictors for alpha and beta power during naturalistic story listening. **A** Example of dependency parsing (Mel’cuk, 1988) and the extracted dependency features (number of opened/ remained open/ resolved dependencies). **B** Model construction. The base model includes low-level linguistic features (acoustic edges, word onset, word frequency), which are included in the dependency models. **C** Schematic of the TRF analysis pipeline. **D** Example of mismatch model construction. The actual feature values are replaced by those of another story, while keeping the initial positions. **E** Grand average power spectrum over all data, participants, and sensors. The FOOOF algorithm (Donoghue et al., 2020) was used to separate the fractal from the oscillatory components of the original signal.

We used an automated parser (Stanford parser “Stanza”; Qi et al., 2020) to generate dependency graphs for each sentence in the stories. Stanza uses Universal Dependencies (Nivre et al., 2016), which is a set of dependency relations that are cross-linguistically applicable (see Table S1 in Supplementary material for the types of Universal Dependencies in our stories). Based on those, three dependency measures were extracted for each word using custom-written scripts: (1) number of opened dependencies, (2) number of dependencies that remained open, and (3) number of resolved dependencies. As we did not have any hypothesis about left-vs. right-branching dependencies, we summed over both directions (Fig. 1A). Dependency features were represented as valued impulses at the word onsets of the respective words where the dependency took place.

1. *Opened dependencies:* the number of dependencies that open at a given word. In the example of Fig. 1A, one dependency opens at each word (nsubj, obj, det, amod, respectively) except for the last one (zero opened dependencies).
2. *Remained open dependencies:* the number of dependencies that are already open but remain unresolved. In Fig. 1A, the obj dependency is still unresolved at word “the”, while both the obj and det dependencies are unresolved at word “big”.
3. *Resolved dependencies:* the number of dependencies that are resolved at this word. In Fig. 1A, the nsubj relation is resolved at word “opened”, the rest of the dependencies are resolved at the last word “presents”.

As mentioned earlier, we were interested in investigating high-level cognitive processing associated with comprehension. Content words (nouns, verbs, adjectives, adverbs) are known to have a considerably specific semantic content and carry the main meaning of the sentence, whereas function words (pronouns, articles, prepositions, auxiliary verbs) contribute mostly to the grammatical structure and have a less conceptual meaning (Corver & van Riemsdijk, 2013). Additionally, function words are usually a part of short-distance relations. Therefore, we focused our analysis on dependency relations between content words only. This was done by analyzing dependencies that were comprised of two content words, while excluding dependencies in which at least one of the two words was a function word.

##### Low-level features: control variables

To make sure that potential dependency effects are not due to low-level linguistic properties, we considered the following features as our base model:

###### Acoustic edges

Abrupt changes in the acoustics are tracked by neural activity. First, we generated broadband envelopes of the audio files using gammatone filter banks (method following Fishbach et al., 2001). Then, we calculated the derivative of the envelope and defined as acoustic edges the points when the derivative exceeds its 97.5th percentile. Acoustic edges were represented as non-zero, equally valued pulses.

###### Word onset

Neural activity has been found to track the onset of words, due to the brain’s parsing of the acoustic input to form discrete meaningful units (Ding & Simon, 2014). Word onsets were represented as non-zero, equally valued pulses at the time points defined by the forced alignment procedure.

###### Word frequency

The frequency of a word outside the sentential context has been shown to modulate neural responses (Brodbeck et al., 2018). Two online databases were used to calculate word frequency, SUBTLEX-NL for Dutch (Keuleers et al., 2010) and Lexique for French (New et al., 2004). The number of instances of each word was divided by the total number of instances of all words. Word frequency was defined as the negative logarithm of that number, so that the higher the value, the lower the frequency.

Before using the above features in the linear regression analysis, we examined the feature inter-correlations by calculating Pearson’s *r* coefficient between features. All correlations were low to moderate (see Fig. S1 of Supplementary material). To detect multicollinearity between features, the *variance inflation factor* (VIF) was computed, which indicates whether the variation of one feature is largely explained by a linear combination of the other features. VIF was low for all features (*NL*_*ac_edges*_ = 1, *NL*_*freq*_ = 1.34, *NL*_*opened*_ = 1.18, *NL*_*remained_open*_ = 1.10, *NL*_*resolved*_ = 1.24; *FR*_*ac_edges*_ = 1, *FR*_*freq*_ = 3.09, *FR*_*opened*_ = 1.44, *FR*_*remained_open*_ = 1.66, *FR*_*resolved*_ = 1.70), indicating no concern for multicollinearity (see Table S2 in Supplementary material for the feature descriptives). Finally, for each linguistic feature, values were standardized to have unit variance and zero mean.

#### 2.5.4. Alpha and beta power estimation

Following our hypotheses regarding alpha and beta oscillations, we used spectral analysis of the MEG data to confirm the presence of two distinct peaks separately for each band, as an index of oscillatory activity. Welch’s method was used to compute the power spectra. Subsequently, the Fitting Oscillations & One Over F (FOOOF) algorithm (Donoghue et al., 2020) was applied to confirm the presence of peaks with power over and above the aperiodic 1/f signal (Fig. 1E).

##### Sensor-level

The time course of the alpha (mean of 8-12 Hz) and beta (15-30 Hz) power was estimated throughout all story parts. Preprocessed MEG data were convolved with a sliding window Hanning taper (adaptive window length). The time-frequency representation was calculated with 1-Hz steps using 6-cycle wavelets over the course of each story part. Then, the wavelet convolved values were averaged over the frequency band.

##### Source-level

First, the complex Fourier coefficients were estimated separately for the alpha and beta bands with same parameters as in the sensor-level analysis. Then, the coefficients were multiplied with the participant’s spatial filter, and, finally, the power (abs^2) of that product was calculated.

Finally, the spectral data were normalized by subtracting the mean alpha and beta power over all time points of all stories from each time point, separately for each sensor/source, before estimation of the TRFs.

#### 2.5.5. Temporal Response Function analysis

We constructed linear forward models (Temporal Response Functions, TRFs: Crosse et al., 2016) to predict alpha and beta power from these dependency features, controlling for low-level linguistic features (Fig. 1C). TRF analysis is capable of disentangling overlapping neural responses due to consecutive events with high temporal proximity, and can handle confounding covariates. TRFs are forward or encoding models based on the assumption that the output of a system relates to the input via a linear convolution (Broderick et al., 2018; Ding & Simon, 2012). Here we assume that the neural responses (alpha and beta power) at each sensor can be expressed as a linear combination of linguistic features shifted by different latencies (see Fig. 1C for a schematic of the TRF analysis pipeline). Specifically, the instantaneous MEG response *r* (*t, n*) of times *t* = 1 …*T* at channel *n* is expressed by the convolution of a linguistic feature, *s*_*k*_ (t), with a kernel or TRF, *w*_*k*_ (*τ, n*). The TRF covers a specified range of time lags, *τ*, relative to time t. The forward model can be represented by the following equation:

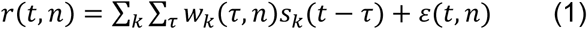

where *ε*(*t*, *n*) is a white noise process, capturing part of the signal unrelated to the stimulus. Note that contributions from each feature are linearly combined. The TRF is estimated by minimizing the mean-squared error between the MEG response, *r* (*t*, *n*), and the predicted MEG response,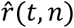

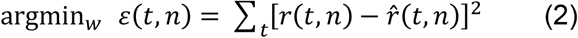

The solution to (2) can be computed in closed-form using the pseudo inverse:

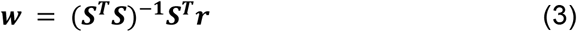

where ***S*** is the concatenation of the lagged time series of each linguistic feature, *s*_*k*_.

TRF analysis was conducted using the MATLAB mTRF Toolbox (Crosse et al., 2016). Here we used the function *mTRFtrain* to estimate the TRF coefficients for each linguistic feature, separately for NL and FR stories. By visual inspection of the TRF coefficients, the time lags over which TRFs were analyzed were from -1 to 1.5 s. The TRF at time *t* indexes how a unit change in a given linguistic feature affects the MEG response *t* seconds later. For Ridge regression, a regularization term is added to leverage the fact that the inversion of ***S***^***T***^***S*** is unstable, and thus prevent overfitting due to fitting high-frequency noise. This happens when the columns of ***S*** are correlated. With continuous regressors, the lagged time series forming the columns of ***S*** comprise a highly autocorrelated signal. However, in our case, all columns of the lagged time series are independent, as they are not continuous and contain nonzero values only at word onsets. The lagged time series is thus not correlated, hence adding a regularization term was not necessary and would lead to underfitting.

##### Model validation

Validation of the TRF models was performed by comparing the Pearson’s *r* correlation between the actual MEG and the reconstructed MEG response. This was implemented using the function *mTRFcrossval* of the mTRF Toolbox following a leave-one-out cross-validation approach. Specifically, a story part was used as the test set and the remaining *M*-1 story parts were used as the training set. The TRF model was then estimated for each story part of the training set, and their average TRF is computed. Subsequently, the averaged model was convolved with the test set to predict the MEG responses. Pearson’s *r* was computed between the actual MEG and the reconstructed MEG responses of the test set from -1 to 1.5 s. The aforementioned process was repeated *M* times, so that all story parts were assigned to the test set once. The Pearson’s *r* values were then averaged over all *M* validations. This procedure was done separately for NL and FR stories.

### 2.6. Statistical evaluation

#### 2.6.1. Model comparison

We first assessed the contribution of the low-level linguistic features to alpha and beta power modulations by evaluating their reconstruction accuracy. Results showed that all features (acoustic edges, word onset, word frequency) explain a substantial amount of variance of the MEG response (see section “Speech features evaluation” in Supplementary material).

##### Comparison with base model

In order to test whether the dependency features predict the neural data over and beyond the base model, we compared reconstruction accuracy between the base model (including only the low-level linguistic features) against the base model augmented with each and all of the dependency features (opened/ remained open/ resolved/ full) (see Fig. 1B for model construction).

As dependencies do not occur at every word instance, dependency features had a substantially lower number of non-zero values (“trials” from now on) compared to the base features (base features: *N*_*word_onset, word_frequency*_ = 8535; dependency features: *N*_*opened*_ = 2691; *N*_*remained_open*_ = 5596; *N*_*resolved*_ = 2381). This would affect the signal-to-noise ratio in the estimated TRF and, therefore, the associated reconstruction accuracy. In order to balance the number of trials between features, a bootstrapping procedure with replacement was followed for model comparison. More specifically, the feature with the smallest number of trials was first identified. Then, an equal number of trials was randomly selected in the rest of the features of the two models being compared at that time, while the excessive trials were converted to zero. This was performed for every feature except acoustic edges, as those trials were not aligned with word onset and were therefore relatively independent to the rest of the features. To make sure that our effects would not be due to a certain random selection during bootstrapping, we repeated this procedure 10,000 times. Subsequently, we tested in how many of these iterations reconstruction accuracy of the dependency model was significantly higher than the base model. To do this, reconstruction accuracy was first averaged over sensors exhibiting improved accuracy over the base model, i.e., where the z-scored difference between models exceeded one standard deviation. Then, paired *t*-tests were conducted between models. Results showed that all models significantly improved reconstruction accuracy compared to the base model across the 10,000 iterations (percentage of significant iterations: > 95%) of the bootstrapping procedure (see “Model comparison” in Supplementary material for the *t* value distributions). Therefore, reconstruction accuracy was averaged over all iterations to perform the final statistical evaluation. Reconstruction accuracy was then averaged over the sensors of which the z-score difference exceeded one standard deviation in more than 50% of the iterations.

We performed a 3 (model: opened/ remained open/ resolved) x 2 (frequency band: alpha vs. beta) repeated-measures ANOVA with reconstruction accuracy improvement (dependency model – base model) as the dependent variable. Improvement was also compared to zero with one-sample *t*-tests, for each model. As results showed that all three dependency features were significant, the full model was evaluated in a 4 (model: opened/ remained open/ resolved/ full) x 2 (frequency band: alpha vs. beta) repeated-measures ANOVA, and was compared to zero. All post hoc contrasts were Bonferroni corrected for multiple comparisons.

##### Comparison with mismatch model

To confirm that reconstruction accuracy improvement with the dependency features is not merely due to (1) the addition of features or (2) the existence or not of a dependency state independent of its value, we compared the full model with mismatch models. Mismatch models are models of which the feature values of one of the features is replaced by those from another story, while keeping the value positions of the actual story (Fig. 1D). This allows to compare models with matching number of predictors.

As the mismatch models have the same number of trials per feature with the actual models, there was no need for a bootstrapping approach here. We performed a 4 (model: opened/ remained open/ resolved/ full) x 2 (frequency band: alpha vs. beta) repeated-measures ANOVA with reconstruction accuracy improvement as the dependent variable, averaged over sensors with z-scored difference between mismatch – actual exceeding 1 *SD*. One-sample *t*-tests compared reconstruction accuracy improvement from zero.

##### Control analysis:Comparison with mismatch model in French stories

As participants did not understand French, we used the French stories as a control condition to confirm that the alpha and beta power modulations by the dependency features in Dutch is linked to comprehension rather than acoustic or speech properties. Similar to the above analysis, we performed one-sample *t*-tests (compared to zero) and a 3 × 2 ANOVA on the reconstruction accuracy difference between actual vs. mismatch models, averaged over the identified sensors with maximal improvement.

#### 2.6.2. Reconstruction accuracy between Dutch (NL) vs. French (FR) stories

Considering the multiple comparisons problem and the lack of a specific hypothesis about the location of the effects, we used a non-parametric cluster permutation approach (Maris & Oostenveld, 2007) to compare the reconstruction accuracy in NL vs. FR on source level. As there were nine story parts in NL, but only four in FR, we performed a bootstrapping procedure with replacement by randomly selecting four of the NL story parts over which we compared the reconstruction accuracy with the FR. This was done over 50 iterations, all showing significant differences between conditions. Reconstruction accuracy was then averaged over all iterations to perform the final statistical evaluation. The cluster permutation procedure addresses the multiple comparison problem by combining neighbouring source points that show the same effect into clusters and comparing those with the null distribution. Paired samples *t* tests were computed for each source point, testing NL vs. FR conditions. Spatially adjacent source points whose *t* values exceeded an a priori threshold (uncorrected *p* value < .05) were combined into the same cluster, with the cluster-level statistic calculated as the sum of the *t* values of the cluster. Finally, the values of the cluster-level statistic were evaluated by calculating the probability that it would be observed under the assumption that the two compared conditions are not significantly different (*α* = .05, two-tailed). To obtain a null distribution to evaluate the statistic of the actual data, values were randomly assigned to the two conditions and the statistics re-computed 1,000 times (Monte-Carlo permutation).

#### 2.6.3. Alpha and beta band modulations by dependency features

##### Comparison of dependency models with base model:*reconstruction accuracy*

To identify the neural sources of the dependency feature contributions, we used beamformer source reconstruction in the alpha and beta bands (see Methods section 4.5.2. and 4.5.4.). The reconstruction accuracy of each dependency model was compared with the base model. A bootstrapping approach with replacement was used as described in section 4.6.1., so that all features of the two models under comparison had the same number of trials. Reconstruction accuracy at each source point was averaged over iterations. A cluster-based permutation approach was used for statistical evaluation as described in section 4.6.2. (*α* = .05, one-tailed; 1,000 permutations), by shuffling the labels between dependency and base model. This analysis was performed separately for the alpha and beta bands.

##### Comparison of dependency features with mismatch model:*TRF coefficients*

In order to define the contribution of the dependency features in time, the TRF coefficients were analyzed. Specifically, the coefficients of each dependency feature were averaged over significant sources, as identified from the aforementioned analysis in section 2.6.3, therefore the comparison was done over the temporal, but not spatial, dimension. This was done both for the full model, as well as for the respective mismatch models of each dependency feature. Subsequently, a cluster-based permutation procedure was performed between the time courses of the averaged coefficients in the actual vs. mismatch models, separately for opened/ remained open/ resolved coefficients and their respective mismatch models (*α* = .05, two-tailed; 2,000 permutations). Note that here clusters were formed along the time dimension (i.e., based on temporal rather than spatial adjacency).

The Brainnetome Atlas (Fan et al., 2016) was used to identify the regions (parcel labels) where the effects were found. According to this atlas, each hemisphere is divided into 123 parcels, while the parcellation is based on both structural and functional connectivity.

##### Data availability

Data and code used in the main analyses are available from the corresponding author on reasonable request.

## 3. Results

### 3.1. Comprehension of Dutch but not French stories

To confirm that participants paid attention to and understood the NL stories but not the FR stories, we calculated performance accuracy as the percentage of correct answers in the multiple-choice comprehension questions. Results revealed that participants comprehended NL stories (*M* = 89.0, *SD* = 6.3) significantly better than FR stories (*M* = 25.7, *SD* = 10.7) (*t*(21) = 23.907, *p* < .001, Cohen’s *d* = 7.189). Performance was significantly higher than chance level (25%) only for NL (*t*(21) = 47.337, *p* < .001, Cohen’s *d* = 10.092), but not for FR (*t*(21) = .298, *p* = .768, Cohen’s *d* = .063) (see “Behavioural data” in Supplementary material).

### 3.2. Dependency features predict alpha and beta power beyond low-level linguistic features

#### 3.2.1. Dependency features improve reconstruction accuracy from base model

To evaluate whether each of the dependency features explains variance of alpha and beta power over and beyond the base model (i.e., based on the features acoustic edges, word onset, word frequency), we compared the reconstruction accuracy (Pearson’s *r*) improvement by adding opened/ remained open/ resolved/ all features to the base model (Fig. 2A). Reconstruction accuracy was averaged over sensors exhibiting improved accuracy over the base model (z-scored improvement > 1 *SD*). A bootstrapping method with replacement was used to ensure equal number of trials between features of each model (see Methods section 4.6.1. for more details on the bootstrapping method).

**Fig. 2.**
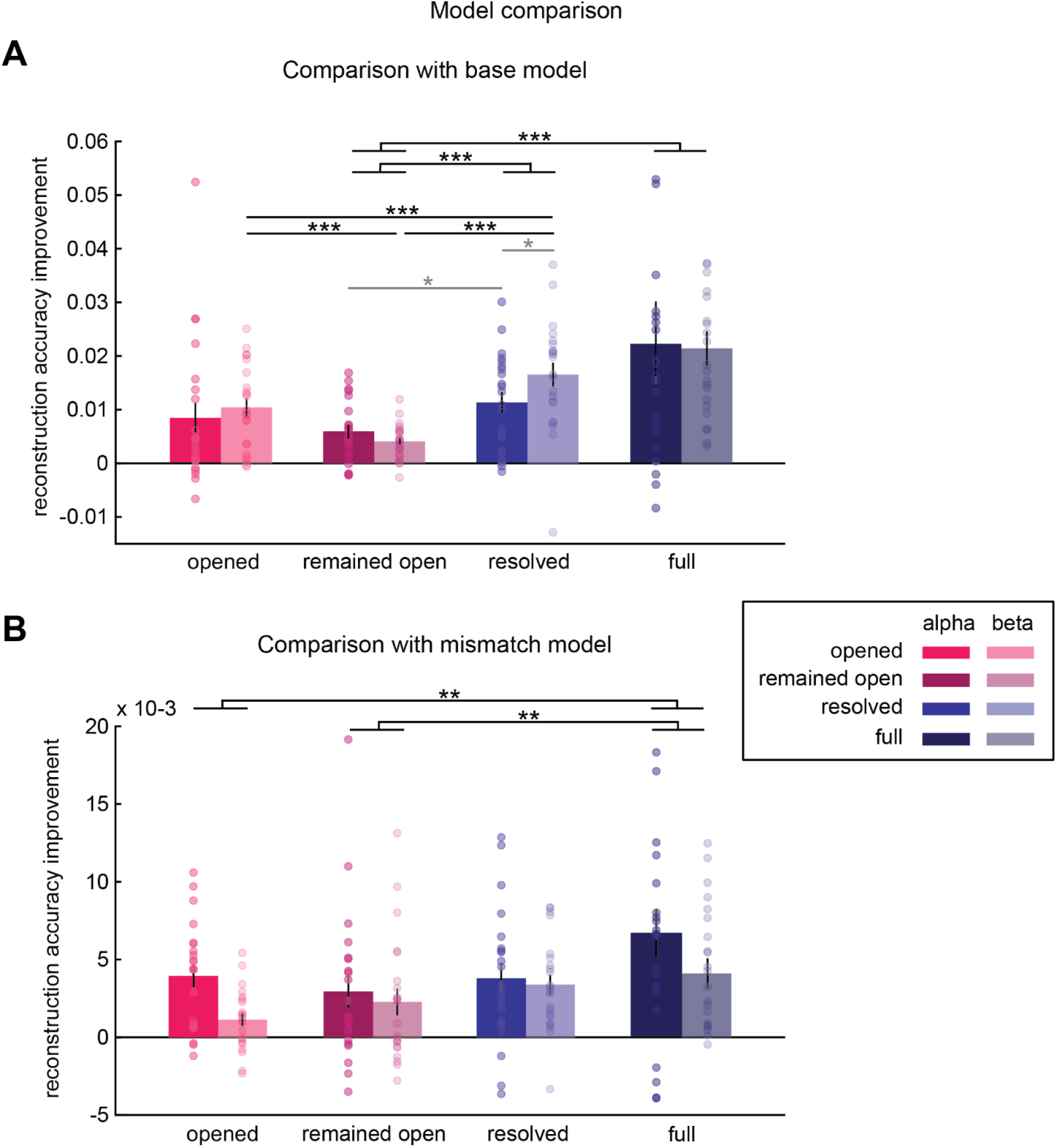
Model comparison: Reconstruction accuracy (Pearson’s *r*) improvement using dependency features. **A** Reconstruction accuracy improvement from base model (acoustic edges, word onset, word frequency) by adding opened/ remained open/ resolved/ all dependency features, averaged over sensors exhibiting improved accuracy over the base model, for alpha (opaque) and beta bands (transparent). **B** Reconstruction accuracy improvement from mismatch models (i.e., models in which the feature values of the respective feature are replaced by those from another story), averaged over sensors exhibiting improved accuracy over the base model, for alpha and beta bands. For instance, the first bar group entitled “opened” refers to the comparison of the full model with a model in which the feature opened is mismatched, i.e., from a different story. The “full” bar group represents the comparison of the full model with a model in which all dependency features come from other stories. Error bars represent +/-1 *SEM*. * *p* < .05 (gray for uncorrected), ** *p* < .01, *** *p* < .001.

Reconstruction accuracy improvement was significantly higher than zero in all models and bands (alpha — opened: *t*(21) = 2.915, *p* = .008; remained open: *t*(21) = 4.903, *p* < .001; resolved: *t*(21) = 5.889, *p* < .001; beta — opened: *t*(21) = 6.205, *p* < .001; remained open: *t*(21) = 5.746, *p* < .001; resolved: *t*(21) = 7.418, *p* < .001).

A 3 (model: opened/ remained open/ resolved) x 2 (band: alpha/ beta) repeated-measures ANOVA revealed a significant *model* x *band* interaction (*F*(2,42) = 3.254, *p* = .049, *η*^*2*^ = .134). Planned contrasts in the beta band were significant: resolved was higher than opened (*t*(21) = 4.465, *p* < .001, Cohen’s *d* = 1.030) and remained open (*t*(21) = 6.170, *p* = .001, Cohen’s *d* = 1.708), and opened was higher than remained open (*t*(21) = 4.001, *p* = .001, Cohen’s *d* = .984). In the alpha band, resolved was higher than remained open, but this did not survive Bonferroni correction at *α* = .005 (*t*(21) = 2.257, *p* = .035, Cohen’s *d* = .491). Resolved was also higher for beta compared to alpha, but this was not significant after multiple comparison correction either (*t*(21) = 2.552, *p* = .019, Cohen’s *d* = .549). None of the other contrasts were significant (*p* > .2).

The ANOVA also revealed a significant effect of *model*, which was due to resolved being higher than remained open (*t*(21) = 4.776, *p* < .001, Cohen’s *d* = 1.151), while there was a trend for resolved being higher than opened (*t*(21) = 2.190, *p* = .040, Cohen’s *d* = .470) and the same for opened compared to remained open (*t*(21) = 2.096, *p* = .048, Cohen’s *d* = .522). There was no significant main effect of frequency band (*p* = .131).

As all three dependency features contributed to explained variance of the alpha and beta power, we created a full model with all features included. A 4 (model: opened/ remained open/ resolved/ full) x 2 (band: alpha/ beta) repeated-measures ANOVA revealed a significant main effect of *model* (*F*(3,63) = 6.004, *p* = .001, *η*^*2*^ = .222), which was due to the full model being higher than the remained open (*t*(21) = 3.269, *p* = .004, Cohen’s *d* = .981) and a trend for the full being higher than the opened (*t*(21) = 2.190, *p* = .040, Cohen’s *d* = .508). The full model was not significantly different from the resolved model (*t*(21) = 1.385, *p* = .181, Cohen’s *d* = .324). There was no main effect of band or interaction between the variables (*p* < .4). Finally, reconstruction accuracy improvement for the full model was significantly higher than zero (alpha: *t*(21) = 2.831, *p* = .010; beta: *t*(21) = 6.710, *p* < .001).

#### 3.2.2. Dependency features improve reconstruction accuracy from mismatch model

To test whether reconstruction accuracy improved merely due to the addition of extra features, we compared the full model with mismatch models (Fig. 2B) (for details see Methods section 4.6.1.). Reconstruction accuracy improvement was significantly higher than zero in all models and bands (alpha — opened: *t*(21) = 5.432, *p* < .001; remained open: *t*(21) = 2.784, *p* = .011; resolved: *t*(21) = 4.074, *p* = .001; full: *t*(21) = 4.340, *p* < .001; beta — opened: *t*(21) = 2.726, *p* = .013; remained open: *t*(21) = 2.688, *p* = .014; resolved: *t*(21) = 5.352, *p* < .001; full: *t*(21) = 4.232, *p* < .001). A 4 (model: opened/ remained open/ resolved/ full) x 2 (band: alpha/ beta) repeated-measures ANOVA revealed a significant main effect of model (*F*(3,63) = 4.514, *p* = .006, *η*^*2*^ = .177). This was due to the full model showing higher improvement than both opened (*t*(21) = 2.797, *p* = .011, Cohen’s *d* = .757) and remained open (*t*(21) = 3.484, *p* = .002, Cohen’s *d* = .634). There was no other significant main effect or interaction between the variables (*p* > .06).

#### 3.2.3. Control analysis: Reconstruction accuracy did not improve in French stories

In order to confirm that the effect of the dependency was due to language comprehension rather than any speech properties, we tested whether reconstruction accuracy was significantly higher in dependency vs. mismatch models in the French stories. Conducting the same analysis as in 2.2.2., but for French, we found that reconstruction accuracy was not significantly higher than zero in any of the three dependency models (*p* > .1). There was no significant main effect or interaction between the variables (*p* > .1; see Fig. S5 in Supplementary material).

#### 3.2.4. Dependency features modulate alpha power peaking in left temporal regions

##### Effect of comprehension:Dutch vs. French stories

First, we wanted to investigate the effect of comprehension by comparing the reconstruction accuracies of the full model (acoustic edges, word onset, word frequency, opened, remained open, resolved dependencies) in NL vs. FR stories. A non-parametric cluster permutation test on source level revealed a significant cluster mostly located in left temporal areas (cluster-corrected *p* = .006) for which NL exhibited higher reconstruction accuracy compared to FR (Fig. 3A). The mean *t* values of the significant cluster as well as the percentage of significant source points (voxels) per parcel (based on an anatomical atlas) are shown in Table 1.

**Table 1.**
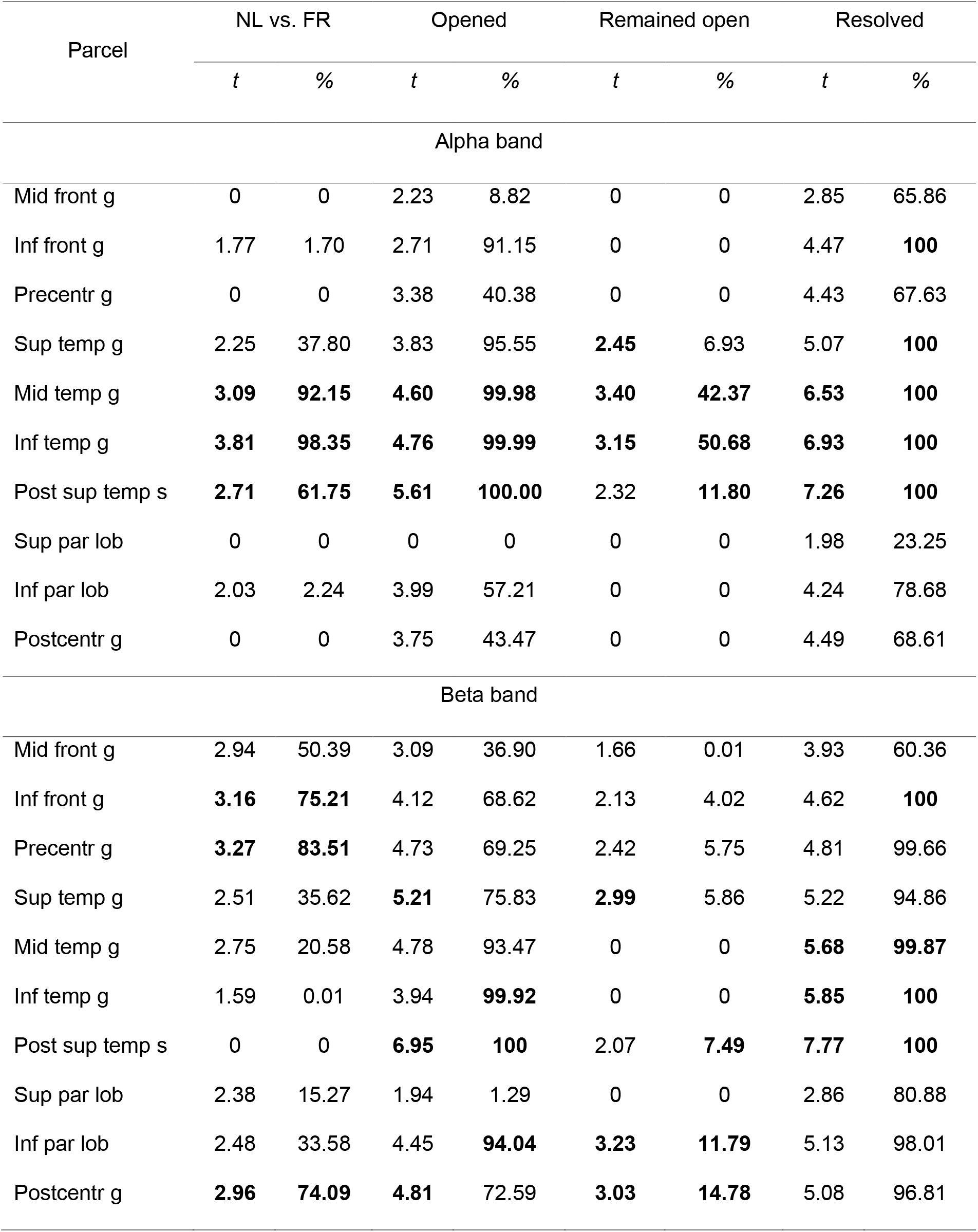
Descriptive statistics of source localization of dependency features effects in the left hemisphere (see Fig. 3). Columns “t” show the mean t value over significant voxels for NL vs. FR and opened/ remained open/ resolved vs. base models, while columns “%” show the percentage of significant voxels in the given parcel. The parcels with values >95% per column are in **bold**.

**Fig. 3.**
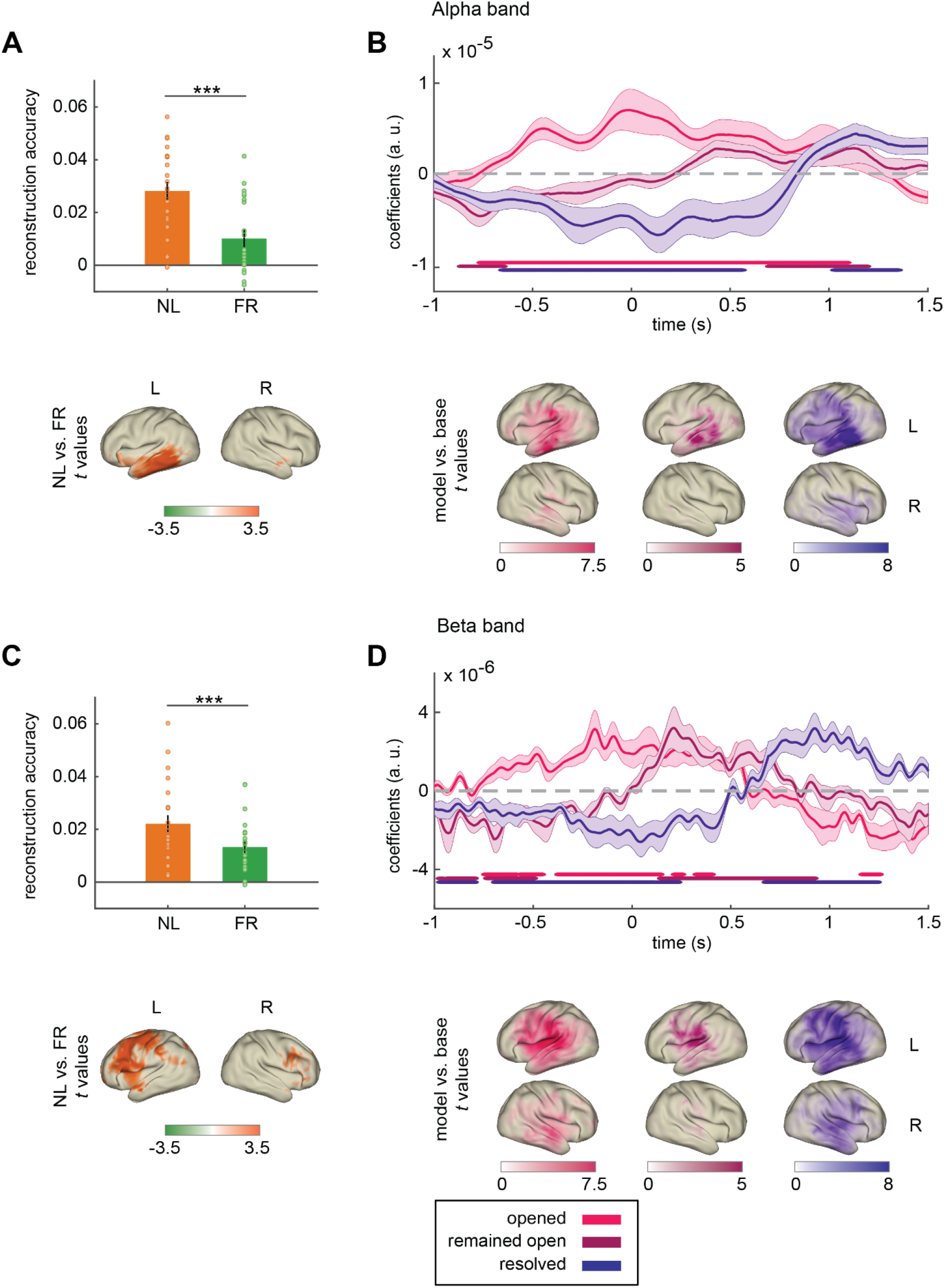
Results of the Temporal Response Function (TRF) analysis separately for the alpha and beta bands. **A** Top: Reconstruction accuracy of the full model in Dutch (NL) vs. French (FR) stories averaged over the significant sources. Bottom: Significant sources for the reconstruction accuracy between NL vs. FR stories based on cluster permutation test. **B** Top: Time courses of the TRF coefficients for each feature. Horizontal lines indicate significant time instances compared to a mismatch model (feature values replaced by those from another story). Bottom: Significant sources for the reconstruction accuracy of models opened/ remained open/ resolved vs. the base model based on a cluster permutation test. **C-D** Same as **A-B**, but for the beta band. Error bars represent +/-1 SEM. *** *p* < .001.

##### Effect of dependency features

In order to identify the contributions of each dependency feature, we performed a cluster permutation test of the reconstruction accuracies in each dependency model (opened/ remained open/ resolved) vs. the base model (Fig. 3B, bottom). We found significant clusters for opened vs. base model (*p* < .001), remained open vs. base (*p* = .022), and, finally, resolved vs. base (*p* < .001). The mean *t* values of the significant clusters as well as the percentage of significant sources per parcel are shown in Table 1 (parcels including cortical areas only, subcortical excluded).

Subsequently, we analyzed the temporal profiles of the aforementioned effects, as demonstrated by their respective TRFs (Fig. 3B, top). More specifically, we averaged over the TRF coefficients of the significant clusters and performed a cluster permutation test in the time dimension between the actual TRF coefficients and those of a mismatch model. With regard to the effect of the opened feature, results revealed a long-lasting positive-going wave in the alpha band from around -.75 to 1.1 s, peaking at word onsets (0 s) (*p* = .002). The remained open feature modulated alpha power negatively around -.75 s (*p* = .002), but positively around .7 to 1.2 s after word onset (*p* = .006). Resolved dependencies showed a long-lasting negativity from around -.7 to .5 s (*p* = .003) and a later positivity around 1 to 1.4 s (*p* = .003).

#### 3.2.5. Dependency features modulate beta power peaking in left frontal, parietal, and temporal regions

##### Effect of comprehensionDutch vs. French stories

We also wanted to investigate the effect of comprehension in the beta band by comparing the reconstruction accuracies of the full model in NL vs. FR stories. A non-parametric cluster permutation test on source level revealed significant clusters, mostly located in left frontal areas (*p* = .002), for which NL exhibited higher reconstruction accuracy compared to FR (Fig. 3C; Table 1).

##### Effect of dependency features

A cluster permutation test of the reconstruction accuracies in each dependency model (opened/ remained open/ resolved) vs. the base model (Fig. 3D, bottom; Table 1) revealed significant clusters for opened vs. base model (*p* < .001), remained open vs. base (*p* = .025), and resolved vs. base (*p* < .001).

Further, we averaged over the TRF coefficients of the aforementioned significant clusters in the beta band and performed a cluster permutation test in the time dimension between the actual TRF coefficients and those of a mismatch model (Fig. 3D, top). Similar to the alpha band, dependency opening was associated with an early positive-going wave in beta band power (*p* = .004), but after around .5 s this became negative (*p* = .004). Remained open started with a short negativity (*p =* .004), and showed a sharp positivity just after word onset, around .25 s, which lasted until 1 s (*p* = .002). Finally, the resolved feature showed an early negativity up until around .25 s (*p* = .003), and a positivity after around .7 s (*p* = .003).

## 4. Discussion

In this study, we tested whether the functional role that alpha and beta oscillations play in low-level perceptual processing can be generalized to naturalistic language processing. Dutch native speakers listened to stories in spoken Dutch and French while MEG was recorded. We identified three states at each word: number of opened/remained open/resolved dependencies. We then constructed forward models to predict alpha and beta power from the dependency features, controlling for low-level linguistic features. In brief, we report the following key findings: (1) High-level syntactic features predict alpha and beta power beyond acoustic, lexical, low-level linguistic features; (2) Left temporal language-related regions are involved in language comprehension for alpha, while frontal and parietal, higher-order language regions, and motor regions play a critical role for beta; and (3) Alpha and beta band dynamics seem to subserve comprehension by contributing to higher-level operations, potentially associated with inhibition and reactivation or propagation processes, during structured meaning composition (Martin, 2020).

As expected, high-level syntactic features predicted alpha and beta power beyond low-level linguistic features. Dependency features improved reconstruction accuracy of a base model with acoustic edges, word onset, and word frequency as predictors. Improvement was not due to the mere addition of predictors, as evidenced by comparisons with mismatch models. Importantly, this was not the case for French, in which the dependency features did not explain more variance in alpha or beta compared to mismatch models. Our results provide evidence for the following: (1) Dependency features are related to alpha and beta modulations in an intelligible but not in an unintelligible language, thus tapping into comprehension processes, and (2) Alpha and beta oscillations are modulated by higher-level operations associated with dependency formation and resolution in naturalistic language processing, beyond low-level features.

Consistent with our first hypothesis, language comprehension seems to involve left temporal areas in the alpha band. Reconstruction accuracy in Dutch stories was significantly higher than French peaking in the middle temporal (MTG), the inferior temporal (ITG), and to a lesser extent the posterior superior temporal gyrus (pSTG). Previous research demonstrated the role of left temporal regions in lexical retrieval and creation of syntactic hierarchies (Klaus et al., 2019; Ouden et al., 2012). The pMTG is argued to receive input from phonological networks and convert sequences of morphemes into nonlinear hierarchical structures, which are then mapped onto semantic networks (Matchin & Hickok, 2020). The pMTG is further thought to be involved in unification operations (Hagoort, 2013; Murphy, 2015). The anterior temporal lobe is associated with syntactic processing (Matchin et al., 2017), though mostly with semantic composition during combinatorial operations (Del Prato & Pylkkänen, 2014; Segaert et al., 2018; Westerlund & Pylkkänen, 2014). An MEG study found that MTG exhibited a high degree of causal outflow to more anterior temporal areas and to the IFG, associated with propagation of information about lexical items to areas performing integration processes during sentence reading (Schoffelen et al., 2017). In our study, dependency resolution was related to integration and unification of the dependent in the sentential context, while dependency formation was responsible for meaning construction during sentence evolution.

On the other hand, linguistic features modulated beta band dynamics in a wider range of regions, peaking in the inferior frontal gyrus (IFG) bilaterally, the left precentral (PrCG) and postcentral (PCG) gyrus, and to a lesser extent the middle frontal gyrus (MFG), with additional contributions from the superior temporal (STG) and middle temporal (MTG) gyrus, and the inferior parietal lobule (IPL). The role of these regions in syntactic hierarchical structure and semantic composition, as well as linguistic unification and integration processes has previously been demonstrated (Berwick et al., 2013; Dronkers et al., 2004; Friederici & von Cramon, 2000; Meyer et al., 2002; Rodd et al., 2005; Zaccarella et al., 2017). The IFG is linked to binding words together into syntactic hierarchies, the integration of more abstract linguistic features with the existing context (Friederici & von Cramon, 2000; Ten Oever et al., 2022; Zaccarella et al., 2017), as well as encoding predictions of syntactic features via top-down evaluation of the fit between an incoming word and the current context (Matchin et al., 2017). Connections originating from temporal areas peak at alpha, whereas connections originating from parietal or frontal regions peak at beta, suggesting frequency-specificity in the language subnetwork (Schoffelen et al., 2017). Finally, the contributions of motor and somatosensory areas might be related to motor-auditory system interactions for efficient speech comprehension (Assaneo & Poeppel, 2018; Keitel et al., 2018; Morillon et al., 2014; Morillon & Baillet, 2017; Poeppel & Assaneo, 2020). For instance, left IFG and PrCG modulate low-frequency activity in left auditory regions during speech comprehension, leading to improved temporal prediction (Abbasi & Gross, 2020; Ten Oever & Martin, 2021; Park et al., 2015; Park et al., 2020; Rimmele et al., 2018; Terporten et al., 2019).

Interestingly, alpha and beta power exhibited a similar time course of activation: an increase before the opening of dependencies, an increase following already opened dependencies, and a decrease prior to dependency resolution, followed by a positive rebound. Despite the lack of substantial temporal dissociation, we observed a distinct spatial pattern of activity. Specifically, alpha and beta power increased before and decreased after the opening of new dependencies, especially beta. The effects were widespread, with peaks in pSTS for both bands, and to a lesser extent in MTG and ITG for alpha, but in STG and PCG for beta. The power increases in language regions reflecting inhibition before dependency opening were surprising; a potential explanation could be reduced interference from already open dependencies (Sauseng et al., 2009; Worden et al., 2000). Further studies are needed to investigate the spatiotemporal dynamics of cortical inhibition during dependency formation. Encoding of new dependencies might be reflected in the alpha and beta desynchronization following the opening of dependencies signaling increased cortical excitability and active processing (Palva & Palva, 2007; Sauseng et al., 2009) or, more speculatively, reflecting coordinate transform from sensory information across the linguistic hierarchy (Martin, 2020).

A power increase was observed immediately following unresolved dependencies for beta, and to a lesser extent for alpha. These effects peaked in temporal regions (STG, MTG, ITG, pSTS) for the alpha band, but in IPL, PCG, and STG for the beta band. Processing difficulty is found to increase with open dependencies (Demberg & Keller, 2008; Vos et al., 2001). Alpha and beta power increases were therefore surprising, as increased load is typically associated with power decreases in task-relevant regions (Jensen & Mazaheri, 2010). On the other hand, there is evidence for higher beta power in long-compared to short-distance dependencies (Meyer et al., 2013), interpreted as active “maintenance” of the current mode of processing (Meyer et al., 2013). It could thus be that beta power increases following unresolved dependencies are linked to ongoing maintenance, but this should be taken with caution, considering that it is still unknown which cognitive operations persist during unresolved dependencies. Future research is needed to study the effects of dependency features on brain responses in a more controlled experimental design, as naturalistic stimuli induce overlapping brain responses sometimes difficult to disentangle.

Finally, we observed alpha and beta power decreases before and a positive rebound following dependency resolution. These effects were widespread but peaked in temporal regions (pSTS, MTG, ITG), while frontal (PrCG, PCG) and parietal (IPL) modulations were stronger in beta than in alpha. Typically, alpha and beta power decreases are associated with upregulation of the cortex, making it more susceptible to the processing of upcoming information (van Ede et al., 2011; Haegens et al., 2011; Schubert et al., 2009) via increased neuronal excitability (Gastaldon et al., 2020; Haegens et al., 2011; Iemi et al., 2022). It could thus be that the observed power decreases reflect the preparation for dependency resolution processing, by a release of language-related regions from inhibition, facilitating processing of upcoming information. The subsequent beta increase is in line with the proposition on beta oscillations supporting content reactivation (Spitzer & Haegens, 2017). According to this view, beta power increases reflect the reactivation of representations by supporting the transition of a latent item to an active working memory state (Antzoulatos & Miller, 2016; Rose et al., 2016). This is based on human and animal studies showing content-specific beta modulations during recall of previously encoded information (Haegens et al., 2011; Haegens et al., 2017; Spitzer et al., 2014; Spitzer & Blankenburg, 2011; Wimmer et al., 2016). Importantly, behavioural work shows that the dependent constituent’s representation is retrieved from memory and reactivated during dependency resolution, in order to be integrated with the current context (Bever & McElree, 1988; Martin & McElree, 2008; McElree et al., 2003; Nicol & Swinney, 1989). We therefore speculate that the observed power increase reflects reactivation of the dependent constituent to achieve successful integration. Overall, our study offers novel contributions for studying alpha and beta dynamics “in the wild” using naturalistic stimuli and TRFs. Finally, our findings provide support for a comprehensive framework regarding the functional role of alpha and beta rhythms, which can be generalized from low-level sensory operations to complex linguistic processes.

## Acknowledgements

We would like to thank Sanne ten Oever for constructive feedback on the study design, and Ryan M.C. Law, Filiz Tezcan Semerci, Cas Coopmans, and Sophie Slaats for contributing to data acquisition. IZ was supported by Big Question 5 of the NWO Gravitation Grant 024.001.006 to the Language in Interaction Consortium. SH was supported by the Netherlands Organisation for Scientific Research (NWO) Vidi grant 016.Vidi.185.137. AEM was supported by an Independent Max Planck Research Group and a Lise Meitner Research Group “Language and Computation in Neural Systems” and by NWO Vidi grant 016.Vidi.188.029.

