## Supplementary material for "Naturalistic language comprehension is supported by alpha and beta oscillations linked to domain-general inhibition and reactivation"

### Methods

#### Universal Dependencies

**Table S1**

*Percentage of trials belonging in each type of Universal Dependency Relation (see* [*https://universaldependencies.org/u/dep/index.html*](https://universaldependencies.org/u/dep/index.html) *for more information) separately for Dutch and French stories. Showing only types occurring >2% of the time.*

| UD type | Dutch | French |
| --- | --- | --- |
| Adverbial modifier | 13.72 | 8.73 |
| Nominal subject | 10.63 | 8.53 |
| Conjunct | 9.25 | 8.03 |
| Oblique nominal | 8.72 | 9.90 |
| Object | 7.21 | 9.23 |
| Verb | 7.04 | 5.47 |
| Determiner | 6.34 | 9.34 |
| Adjectival modifier | 5.41 | 5.88 |
| Case marking | 4.78 | 6.19 |
| Coordinating conjunction | 4.12 | 3.34 |
| Parataxis | 3.00 | 1.04 |
| Nominal modifier | 2.81 | 5.84 |
| Adverbial clause modifier | 2.76 | 2.63 |
| Open clausal complement | 2.73 | 4.08 |
| Marker | 2.58 | 2.04 |
| Auxiliary | 2.09 | 1.45 |

#### Feature descriptives

**Table S2**

*Mean, standard deviation, and range of word frequency, opened, remained open, and resolved features. Acoustic edges and word onset are not presented as they have a single value.*

|  | Dutch | | | French | | |
| --- | --- | --- | --- | --- | --- | --- |
| Feature | *M* | *SD* | *Range* | *M* | *SD* | *Range* |
| Word frequency | 3.13 | 1.39 | 1.39-7.64 | 3.36 | 1.37 | 1.45-8.26 |
| Opened | 1.29 | 0.66 | 1-5 | 1.33 | 0.62 | 1-5 |
| Remained open | 1.70 | 0.86 | 1-7 | 1.53 | 0.74 | 1-5 |
| Resolved | 1.46 | 0.79 | 1-6 | 1.20 | 0.47 | 1-4 |

#### Feature correlations

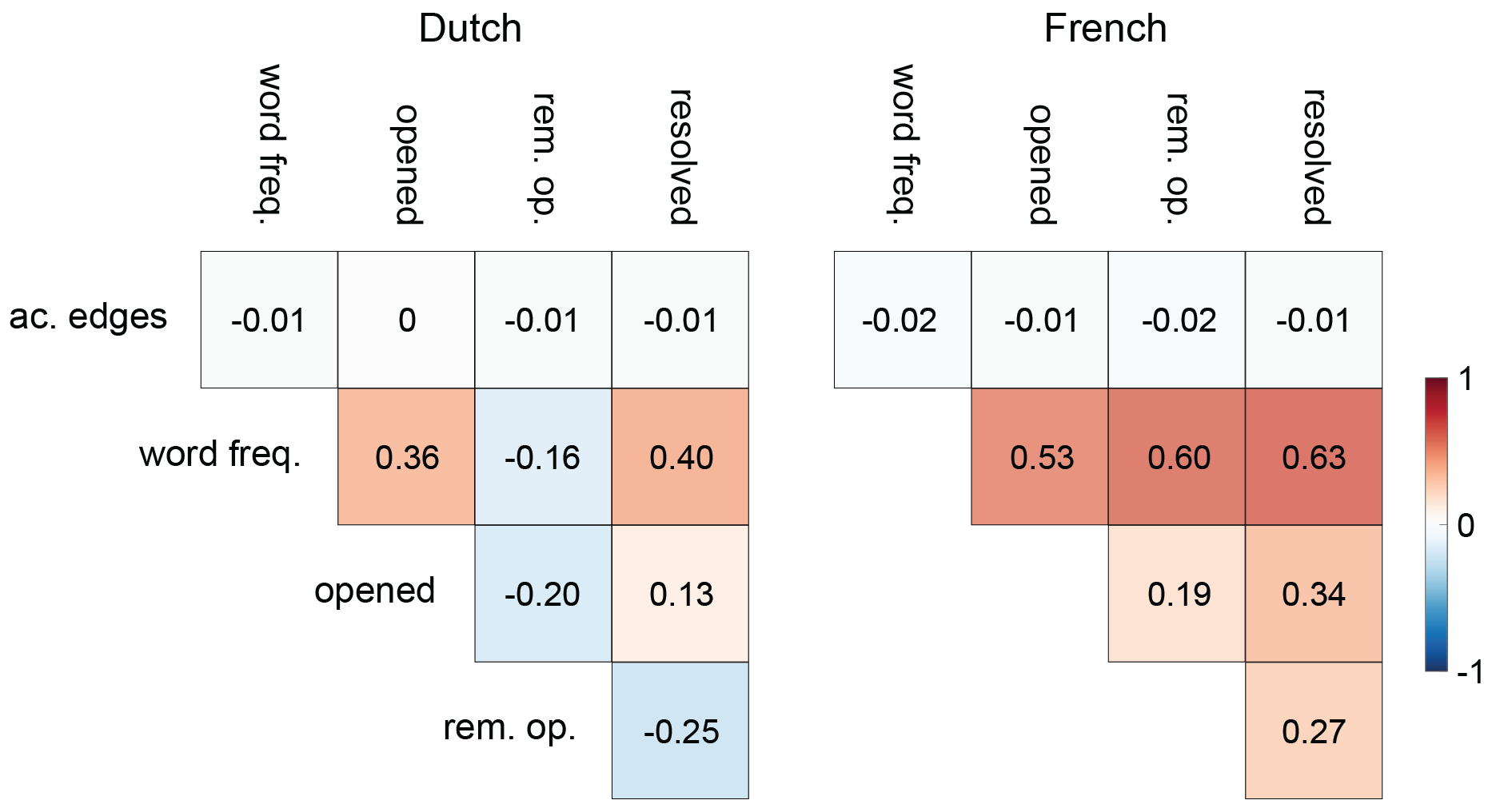

**Fig. S1** Correlation plots between all features (except word onset as it is a constant) separately for Dutch and French stories.

#### Model comparison

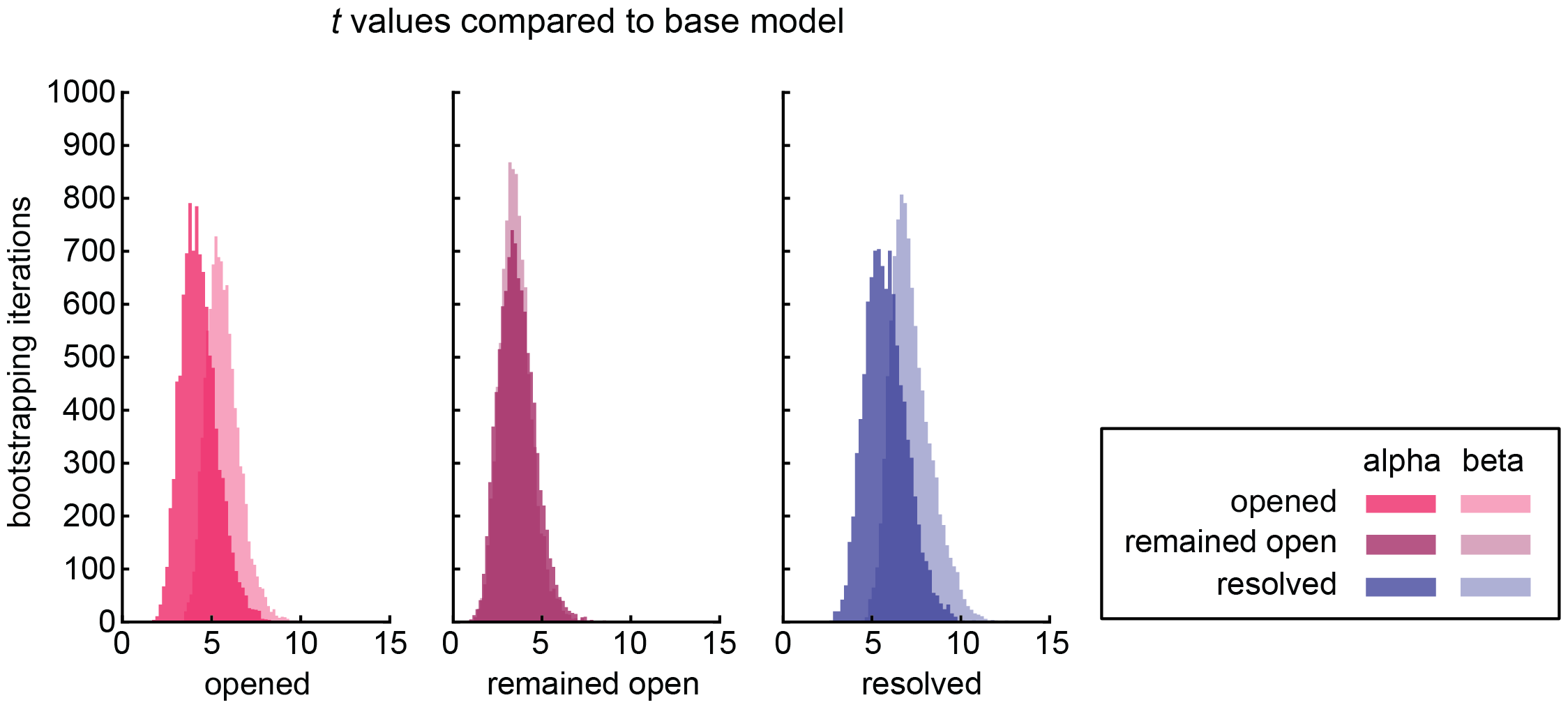

**Fig. S2** Histograms of *t* values between the reconstruction accuracy (Pearson’s *r*) of the models opened/ remained open/ resolved vs. the base model (rec. acc. averaged over sensors exhibiting a z-score difference of > 1 *SD*) for each of the 10000 bootstrapping random trial-selection iterations (see Methods section 2.5.4.1. for details on the bootstrapping procedure).

### Results

#### Behavioural data

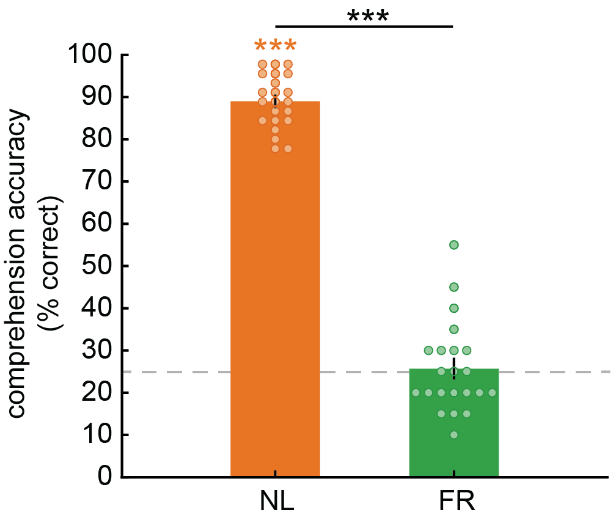

**Fig. S3** Comprehension accuracy defined as percentage of correct responses in the multiple-choice questions for Dutch (NL) vs. French (FR) stories. Coloured asterisks represent significance from zero. Error bars represent +/-1 SEM. *** *p* < .001.

#### Speech features evaluation

We assessed the effect of the speech features by comparing the reconstruction accuracy of a model with acoustic edges vs. adding word onset vs. adding word frequency features (Fig. S3A). Reconstruction accuracy was averaged over sensors >1 *SD*. A 3 (model: acoustic edges/ word onset/ word frequency) x 2 (band: alpha/ beta) x 2 (language: NL/ FR) repeated-measures ANOVA revealed a significant *model* x *band* x *language* interaction (*F*(2,42) = 3.322, *p* = .046, *η^2^* = .137), as well as main effects for *model* (*F*(2,42) = 64.263, *p* < .001, *η^2^* = .754), *band* (*F*(1,21) = 37.377, *p* < .001, *η^2^* = .640), and *language* (*F*(1,21) = 25.306, *p* < .001, *η^2^* = .546). All planned contrasts were significant (*p* < .005), except word onset vs. word frequency in alpha in NL, acoustic edges/ word onset vs. word frequency in alpha in FR, word onset vs. word frequency in beta in FR, and acoustic edges in beta in NL vs. FR (*p* > .07) (Fig. S3B).

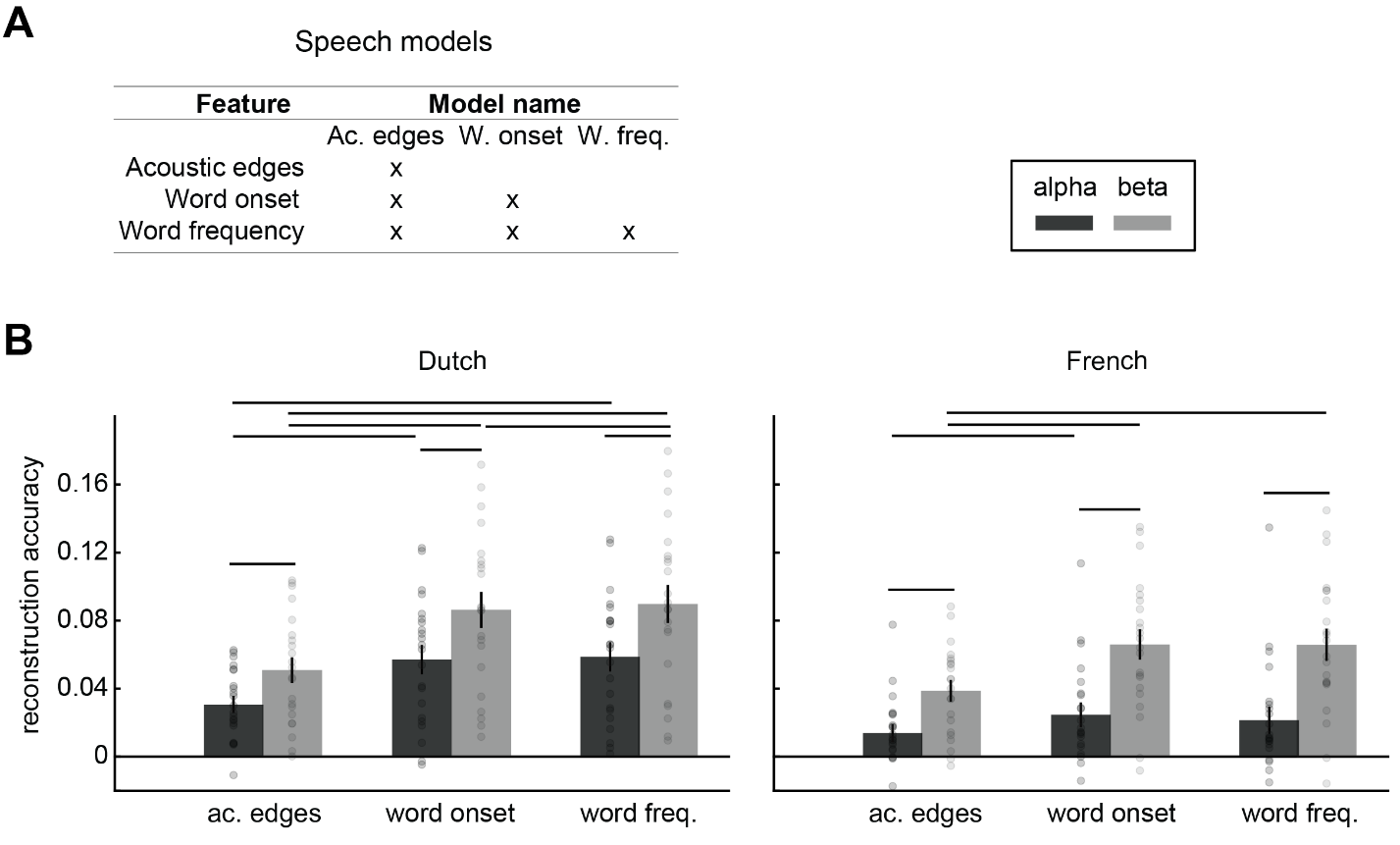

**Fig. S4** Evaluation of the effect of speech features. **A** Speech feature models; **B** Reconstruction accuracy of each model (averaged over sensors >1 *SD*) separately for the alpha (black) and beta (gray) bands, and for Dutch (left) and French (right) stories. Error bars represent +/-1 SEM. Horizontal lines denote statistical significance (*p* < .005).

#### French: Comparison with mismatch models

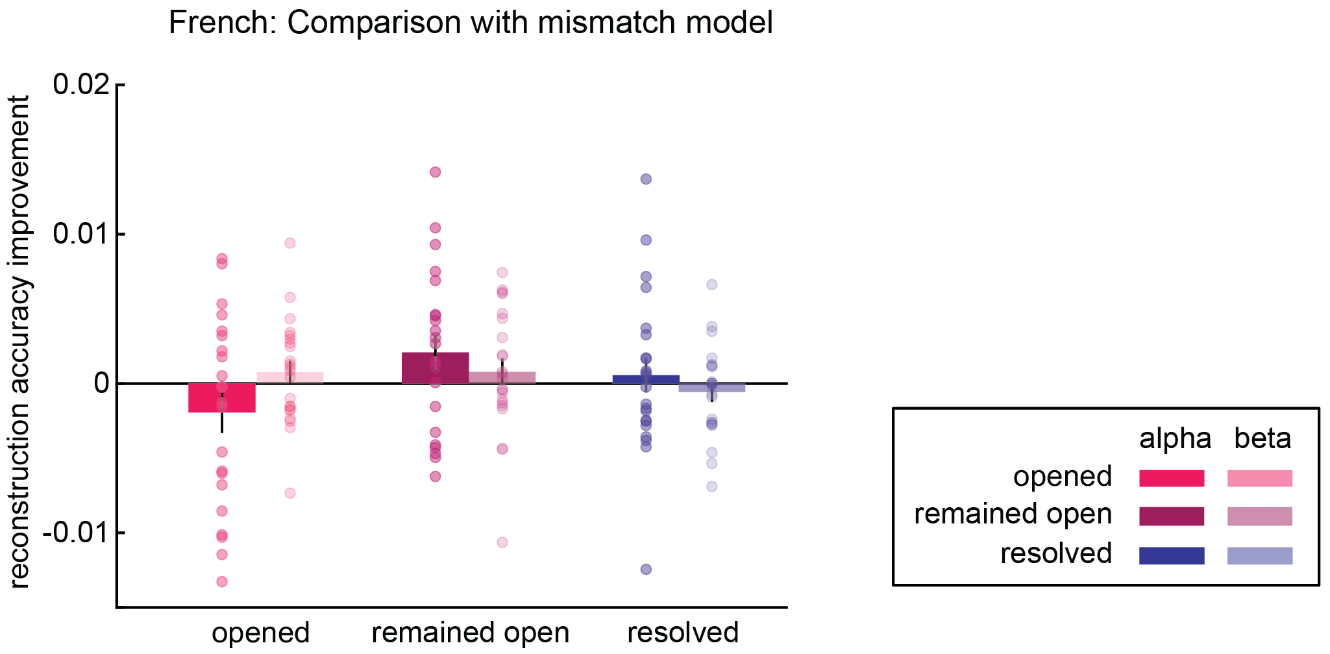

**Fig. S5** Reconstruction accuracy (Pearson’s *r*) improvement from mismatch models (i.e. models in which the feature values of the respective feature are replaced by those from another story) in French stories, for alpha (opaque) and beta (transparent) bands. Error bars represent +/-1 *SEM*.
